# GBZ File Format for Pangenome Graphs

**DOI:** 10.1101/2022.07.12.499787

**Authors:** Jouni Sirén, Benedict Paten

## Abstract

**Motivation:** Pangenome graphs representing aligned genome assemblies are being shared in the text-based Graphical Fragment Assembly format. As the number of assemblies grows, there is a need for a file format that can store the highly repetitive data space-efficiently.

**Results:** We propose the GBZ file format based on data structures used in the Giraffe short read aligner. The format provides good compression, and the files can be efficiently loaded into in-memory data structures. We provide compression and decompression tools and libraries for using GBZ graphs, and we show that they can be efficiently used on a variety of systems.

**Availability:** C++ and Rust implementations are available at https://github.com/jltsiren/gbwtgraph and https://github.com/jltsiren/gbwt-rs, respectively.

**Supplementary information:** Supplementary data are available online.

## 1 Introduction

Pangenome graphs (or genome graphs) (Eizenga *et al*., 2020b) are graphs representing sequence variation. Each node in the graph is labeled with a sequence, and each path represents the concatenation of the labels on it. Pangenome graphs are often used as reference genomes. Initial efforts have focused on using graphs as technical artifacts to improve accuracy in applications such as read alignment (Garrison *et al*., 2018; Kim *et al*., 2019; Rautiainen and Marschall, 2020; Li *et al*., 2020; Sirén *et al*., 2021) and genotyping (Eggertsson *et al*., 2017; Kim *et al*., 2019; Chen *et al*., 2019; Hickey *et al*., 2020; Sirén *et al*., 2021; Ebler *et al*., 2022).

There is increasing interest in using graphs as biologically meaningful references that represent sequence variation in the relevant population. The *Human Pangenome Reference Consortium* (HPRC) (Wang *et al*., 2022) aims to create a human reference genome based on over 350 high quality haplotype-resolved assemblies. The reference will consist of the assembled genomes and pangenome graphs representing their alignments.

Software tools for pangenome graphs use the text-based *Graphical Fragment Assembly* (GFA) format as their data interchange format. GFA was originally intended for assembly graphs, but it is also viable for pangenome graphs with some restrictions and extensions. A GFA file representing only the graph itself is not too large, and it is often compressed further with standard data compressors such as gzip. The format becomes inadequate if we also want to represent the original sequences as paths. While the paths are often highly similar and should compress well, common general-purpose data compressors use windows no larger than a few megabytes. With paths longer than that, the compressor never sees the same region on multiple paths in the same window. Hence it cannot compress the similarities. Decompressing a large text file and reading the graph and the paths into in-memory data structures can also be expensive.

We propose the GBZ file format for pangenome graphs representing aligned sequences. The format is based on the data structures used in the Giraffe aligner (Sirén *et al*., 2021), and it uses a GBWT index (Sirén *et al*., 2020) for storing a large collection of similar paths space-efficiently. GBZ is a path-based format. Paths representing the original sequences are the primary objects, while the existence of nodes and edges is inferred from usage. We provide a standalone C++ library and compressor for creating and using GBZ graphs, as well as a Rust library for using existing graphs. Both libraries can load GBZ graphs quickly into versatile data structures that are space-efficient enough to enable working with human pangenome graphs on a desktop or a laptop.

Our libraries use Elias–Fano encoded bitvectors for storing increasing integer sequences. We describe various improvements to their semantics and the query interface that make them more practical and may be of independent interest.

Pangenome graph construction pipelines sometimes create highly collapsed regions that break the assumptions the GBWT makes to ensure fast queries. We show that decompressing and caching parts of the GBWT index is enough to restore performance in specific tasks when such regions are present. Based on this, we suggest changes to make the in-memory GBWT data structures more robust with highly collapsed graph regions.

## 2 Methods

### 2.1 Background

#### 2.1.1 Strings

In the following, we refer to semiopen and closed integer intervals as [*a* … *b*) = [*a, b*) ∩ ℕ and [*a* … *b*] = [*a, b*] ∩ ℕ, respectively.

A *string S* = *s*_0_ · · · *s_n_*_−1_ of length |*S*| = *n* is a sequence of *characters* from an ordered *alphabet* Σ. For convenience, we often assume that the alphabet is the set of integers [0 … |Σ|). We write *S*[*i*] = *s_i_* and *S*[*i* … *j*) = *s_i_* · · · *s_j_*_−1_ to access individual characters and *substrings* of any sequence *S*. *Prefixes* and *suffixes* are substrings of the form *S*[0 … *j*) and *S*[*i* … *n*), respectively, and we write them as *S*[… *j*) and *S*[*i* …). A *text* string *T* ends with an *endmarker T* [|*T* | − 1] = $ that does not occur in any other text position. The endmarker is the smallest character in the alphabet, and we often assume that $ = 0.

We say that alphabet Σ supports *complementation* if each character *c* ∈ Σ has a *complement* 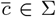 such that 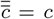 for all *c* ∈ Σ. Given an alphabet Σ that supports complementation, the *reverse complement* of string *S* ∈ Σ^*n*^ is string 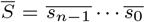 that contains the complement of each character in *S* in reverse order.

Space-efficient data structures rely on *rank* and *select* queries on various sequences. For any string *S*, position *i* ≤ |*S*|, and character *c* ∈ Σ, we define *S*.rank(*i, c*) as the number of occurrences of character *c* in the prefix *S*[… *i*). If *S*[*i*] = *c*, we refer to it as the occurrence of rank *S*.rank(*i, c*) of character *c*. We also define the inverse operation *S*.select(*i, c*) = *j*, where *S*[*j*] is the occurrence of rank *i* of character *c*. (Note that we use 0-based select, where the first occurrence is of rank 0.)

#### 2.1.2 Bitvectors

A *bitvector* is a data structure that encodes a binary sequence (a string over alphabet {0, 1}) and supports efficient rank/select queries. We call characters 0 and 1 *unset* and *set* bits, respectively. If *A* is a strictly increasing sequence of *m* non-negative integers, we can encode it as a bitvector *B*, where *B*[*A*[*i*]] = 1 for all *i < m* and *B*[*j*] = 0 otherwise. If *B* is a binary sequence, we can decode it as integer sequence *A* with *A*[*i*] = *B*.select(*i*, 1). Query *B*.rank(*j*, 1) can then be understood as the number of values *A*[*i*] < *j* in sequence *A*.

*Plain* bitvectors store the binary sequence without any further encoding. There are many efficient structures that support rank and select on plain bitvectors. We use the ones from SDSL (Gog and Petri, 2014).

A binary sequence is *sparse* if the number of set bits is much smaller than the number of unset bits. If a binary sequence is sparse, |*A*| ≪ |*B*| in the corresponding bitvector. In this paper, a sparse bitvector is a data structure that uses *Elias–Fano* encoding to store a sparse binary sequence space-efficiently (Okanohara and Sadakane, 2007).

Elias–Fano encoding stores the low and high parts of each value *A*[*i*] separately. The lowest *w* bits are stored in an integer array low by setting low[*i*] = *A*[*i*] mod 2^*w*^. For the high part, we assign each value *x* to a bucket using bucket(*x*) = ⌊*x*/2^*w*^⌋ and encode the buckets in unary. If a bucket contains *k* values, we encode it as a binary sequence 1^*k*^0. We concatenate the binary sequences for each bucket to form bitvector high. The encoding works best with *w* ≈ log_2_|*B*| − log_2_|*A*|, which makes the number of buckets similar to the number of values. Bitvector high then contains similar numbers of set and unset bits.

If high[*j*] is the set bit of rank *i*, we can compute

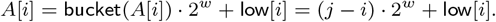

Given *i*, we can compute *j* = high.select(*i*, 1). Hence

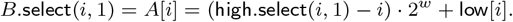

For *B*.rank(*i*, 1), we first find the end of the bucket that would contain value *i* using *h* = high.select(bucket(*i*), 0). The number of values in buckets up to bucket(*i*) is *l* = *h* − bucket(*i*). If high[*h* − 1] = 1, it is the set bit of rank *l* − 1. If also low[*l* − 1] ≥ *i* mod 2^*w*^, we know that *A*[*l* − 1] ≥ *i* and iterate with *h* ← *h* − 1 and *l* ← *l* − 1. If either of the conditions fails, we return *B*.rank(*i*, 1) = *l*.

#### 2.1.3 Burrows–Wheeler transform

Let *T* be a text string of length *n*. The *suffix array* of text *T* is an array SA of pointers to the suffixes of *T* in lexicographic order. For any *i < j*, we have *T* [SA[*i*]…) < *T* [SA[*j*]…). Because the text ends with an endmarker, we always have SA[0] = *n* − 1.

We can generalize the suffix array for an ordered collection *T*_0_*, …, T_m_*_−1_ of *m* texts. Each element of the suffix array is now a pair SA[*i*] = (*j, j*′) that refers to suffix *T_j_* [*j*′…) of text *T_j_*. To make the lexicographic order unique, we assume that the endmarker of *T_i_* is smaller than that of *T_j_* for any *i < j*. Then SA[*i*] = (*i*, |*T_i_*| − 1) for 0 ≤ *i < m*. We use the suffix array to define the *document array* DA as an array such that DA[*i*] = *j* if SA[*i*] refers to a suffix of *T_j_*.

The (multi-string) *Burrows–Wheeler transform* (BWT) (Burrows and Wheeler, 1994) of a collection of texts is a permutation BWT of character occurrences in the texts. For each suffix in lexicographic order, the permutation lists the character preceding the suffix. We define the BWT using the suffix array: BWT[*i*] = *T_j_* [*j*′ − 1] if SA[*i*] = (*j, j*′) and *j*′ > 0. If *j*′ = 0, we set BWT[*i*] = $.

Let C be an array of length |Σ| + 1 such that C[*c*] is the total number of occurrences of all characters *c*′ < *c* in the texts. For any character *c*, if C[*c*] ≤ *i* < C[*c* + 1], and SA[*i*] = (*j, j*′), we know that *T_j_* [*j*′] = *c*. We later use this property for partitioning the BWT into substrings BWT*c* = BWT[C[*c*]… C[*c* + 1]) by the first character of the corresponding suffix.

Given a text collection *T*_0_*, …, T_m_*_−1_, the *lexicographic rank* of string *X* is the number of suffixes of the collection that are smaller than *X*. The key operation on the BWT is the *LF-mapping* that maps the lexicographic rank of a suffix to the lexicographic rank of the preceding suffix. In other words, it is a function such that if SA[*i*] = (*j, j*′), then SA[LF(*i*)] = (*j, j*′ − 1). We leave this form of LF-mapping undefined when *j*′ = 0. A more general form of LF-mapping takes a character *c* ∈ Σ as a second argument. If the lexicographic rank of string *X* is *i*, the lexicographic rank of string *cX* is LF(*i, c*). In particular, LF(*i*) = LF(*i*, BWT[*i*]). We leave the function undefined with *c* = $.

The *FM-index* (Ferragina and Manzini, 2005) is a space-efficient text index based on the BWT. It computes LF-mapping as

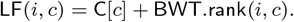

Let SA[*i* … *j*) be the range of suffixes starting with *pattern* (string) *P*. Then the range of suffixes starting with pattern *cP*, for any character *c* ∈ Σ, is SA[LF(*i, c*)… LF(*j, c*)). We can also use LF-mapping for extracting individual texts from the BWT. The last character of text *T_i_* is BWT[*i*]. As long as BWT[*j*] ≠ $ at the current position *j*, there may be other characters in the text before the ones we have already extracted. We therefore move to *j* ← LF(*j*) and find the possible previous character at BWT[*j*].

#### 2.1.4 Graphs

Let *G*′ = (*V*′, *E*′) be a *directed graph* with a finite set of *nodes V*′ ⊂ ℕ and a set of *edges E*′ ⊆ *V*′ × *V*′. Because we often use value 0 for technical purposes and store some information for all integers between node identifiers, we assume that *V*′ is a dense subset of interval [*a* … *b*) for some 1 ≤ *a* < *b*. Let *P* be a string over alphabet *V*′. We say that *P* is a *path* in graph *G*′ if (*P*[*i*], *P*[*i* + 1]) ∈ *E*′ for all 0 ≤ *i* < |*P* | − 1.

Genome graphs are often represented as *bidirected sequence graphs G* = (*V, E, ℓ*). Each node *v* ∈ *V* has a non-empty string *label ℓ*(*v*) over an alphabet that supports complementation, and it can be seen in two *orientations o* ∈ {+, −}. If *o* is an orientation, we use 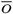 to refer to the other orientation. When we *visit* node *v* in forward orientation +, we read string *ℓ*(*v*, +) = *ℓ*(*v*). For a visit in reverse orientation −, we read the reverse complement 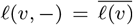. The set of edges *E* is a subset of pairs of node visits (*V* × {+, −}) × (*V* × {+, −}) such that ((*v, o*), (*w, o*′)) ∈ *E* iff 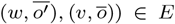. Sequence *P* of node visits is a path in graph *G* if (*P*[*i*], *P*[*i* + 1]) ∈ *E* for all *i*. If *P* is a path in graph *G*, the reverse path is the reverse complement 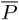, where the complement of a node visit is 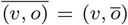. That is, the reverse path visits the same nodes in reverse order and in the other orientation.

Graph representations often store the topology of a bidirected sequence graph *G* = (*V, E, ℓ*) as a directed graph *G*′ = (*V*′, *E*′). Each node visit (*v, o*) ∈ *V* × {+, −} becomes a node *w* ∈ *V*′ in the directed graph.

The *libhandlegraph* library (Eizenga *et al*., 2020a) provides a common C++ interface for various bidirected sequence graph implementations. A *HandleGraph* is an immutable graph. *PathHandleGraph* extends it with a set of named paths, where each path has a unique non-empty string name.

#### 2.1.5 GFA file format

We restrict our attention to the subset of the GFA file format version 1.1 used by HPRC.^1^ As GBZ is not intended for assembly graphs, we do not support features like sequence overlaps between adjacent segments or read coverage annotations. The subset is mostly compatible with the bidirected sequence graph data model:

- *S-lines* or *segments* are the nodes of the graph. They consist of a unique non-empty string name and a non-empty string label.
- *L-lines* or *links* are the edges of the graph. They connect two segment visits. We do not allow any overlaps between the connected segments.
- *P-lines* or *paths* are named paths in the graph. They consist of a unique non-empty string name and a non-empty sequence of segment visits.
- *W-lines* or *walks* are paths with metadata suitable for representing assembled genomes. They consist of a sample name (non-empty string), haplotype identifier (integer), sequence name (non-empty string), an optional interval of positions in the named sequence, and a non-empty sequence of segment visits.

#### 2.1.6 GBWT index

The *GBWT* (Sirén *et al*., 2020) is a run-length encoded FM-index that stores a collection of paths in directed graph *G*′ = (*V*′, *E*′) as strings over alphabet *V*′. Run-length encoding allows it to compress collections of similar paths well. Because we build the FM-index for reverse strings, LF-mapping with character *w* after character *v* follows edge (*v, w*) instead of going backwards on edge (*w, v*). We break the BWT into substrings BWT*v* for each *v* ∈ *V*′ and store the substrings in the nodes. Each node *v* ∈ *V*′ also stores a list of outgoing edges (*v, w*) and some rank information that allows computing LF-mapping using locally stored information. We compress the nodes as byte sequences. GBWT queries involve decompressing entire nodes. Hence we assume that nodes do not have too many neighbors and paths do not visit them too many times.

We use the GBWT for storing paths in a bidirected sequence graph *G* = (*V, E, ℓ*) with the approach described in Section 2.1.4. The GBWT index is usually *bidirectional* (Lam *et al*., 2009). For each path *P*, we store both *P* and 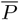 in the index, which enables extending the pattern in both directions.

Each path in the graph (which may be stored as two paths in the GBWT index) may be associated with a structured name. The name consists of four integer fields: sample identifier, contig identifier, haplotype/phase identifier, and fragment identifier / running count. Each name is assumed to be unique. GBWT metadata may also contain lists of string names corresponding to sample and contig identifiers.

The *GBWTGraph* (Sirén *et al*., 2021) is the graph representation used in the Giraffe aligner. It uses a bidirectional GBWT index for graph topology and stores the labels of both orientations of all nodes as a single concatenated string. The graph also caches some information that can be computed from the GBWT for faster access. A GBWTGraph is always a subgraph induced by the paths. Paths representing DNA sequences are the primary objects, while the existence of nodes and edges is inferred from usage. This is not a fundamental limitation of the data structures but a deliberate choice made due to the path-centric algorithms used in Giraffe.

### 2.2 Improved sparse bitvectors

Elias–Fano encoded sparse bitvectors are often used for partitioning integer intervals [*a* … *b*) into subintervals [select(*i*, 1)… select(*i* + 1, 1)). For example, we may want to store an ordered collection of strings *S*_0_, …, *S_m_*_−1_ as a single concatenated string *X* for faster disk I/O. A sparse bitvector *B* of length |*X*| can then serve as an index. If we set *B*[*j*] = 1 at the starting position of each string, we can access the original strings as *S_i_* = *X*[*B*.select(*i*, 1) … *B*.select(*i* + 1, 1)) (with *B*.select(*m*, 1) = |*X*| for convenience). However, bitvector semantics and the rank/select query interface limit the practicality of this approach.

If some of the strings *S*[*i*] are empty, we have *A*[*i*] = *A*[*i* + 1], which would require setting bit *B*[*A*[*i*]] twice. However, Elias–Fano encoding is not limited to storing strictly increasing sequences. The encoding and select(·, 1) queries work correctly as long as all values in bucket *i* occur before the values in bucket *j* in the integer sequence *A*, for all buckets *i < j*. For rank(·, 1) queries, the algorithm works correctly as long as the sequence is non-decreasing. The semantics of rank(·, 0) and select(·, 0) queries become unclear when duplicate values are allowed.

When we want to retrieve string *S*[*i*], we have to compute both *B*.select(*i*, 1) and *B*.select(*i* + 1, 1). Computing them directly is inefficient, as they involve relatively expensive high.select(·, 1) queries. To avoid this, select queries should return an iterator with internal state (*i*, high.select(*i*, 1)). We can compute *B*.select(*i*, 1) efficiently from those values. When we want to advance the iterator, we set *i* ← *i* + 1 and scan for the position of the next set bit in high. This is efficient on the average, as we have chosen a width *w* such that high contains similar numbers of set and unset bits. The iterator can also be used for decompressing the integer sequence *A* efficiently.

Given a value *j* ∈ [*a* … *b*), we may want to retrieve the subinterval [select(*i*, 1)… select(*i* + 1, 1)) containing it, as well as the rank *i* of the subinterval. This can be achieved using a *predecessor* query *B*.pred(*j*) = (*i, B*.select(*i*, 1)), where *i* is the last position with *A*[*i*] ≤ *j*. We can compute it naively by first finding the rank *i* = *B*.rank(*j* + 1, 1) − 1 and then computing *B*.select(*i*, 1). This is inefficient, as rank queries on *B* involve computing high.select(·, 0), while select queries on *B* require high.select(·, 1). A better implementation is similar to a rank query. We first find the end of the bucket that would contain value *j*, and then we iterate backward until we find a value *A*[*i*] ≤ *j* or run out of values.

### 2.3 GBZ file format

#### 2.3.1 General

The GBZ file format is based on the serialization format used in the Simple-SDS library. A file is an array of *elements*: unsigned little-endian 64-bit integers. If the file is memory-mapped, all objects with alignment of 8 bytes or less are properly aligned in memory. A *vector* is serialized by writing the number of items as an element, followed by the concatenated items. If the size of an item is not a multiple of 64 bits, the vector must be padded with 0-bits. Optional or implementation-dependent structures may be serialized by writing the size of the serialized object in elements as an element, followed by the object itself. A reader can then easily skip the object or pass it through as a vector of elements.

Two general principles have guided the development of the GBZ file format: determinism and backward compatibility with the in-memory representation of existing GBWT indexes. The former means that the same data should always be compressed in the same way, without any variation that depends on the implementation or the execution environment. When these principles are in conflict, compatibility has prevailed in the first version of the file format. Compatibility also required making some complex requirements to support old GBWT indexes. Future versions of the file format may choose simplicity and determinism over compatibility.

See Figure 1 for an overview of the file format and Supplement 1 for further details.

**Fig. 1.**
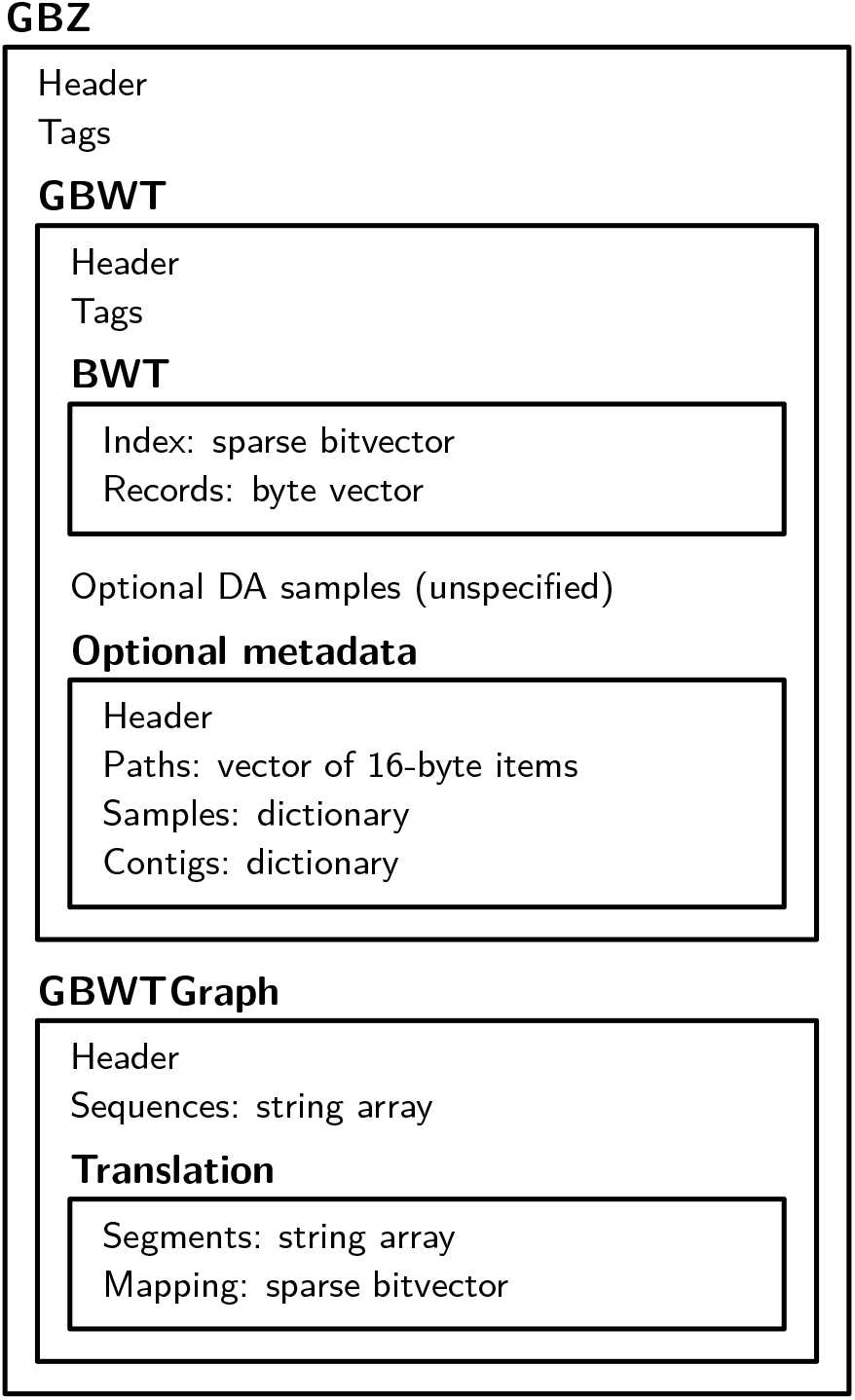
Overview of the GBZ file format.

#### 2.3.2 Building blocks

Simple-SDS provides a number of basic succinct data structures. They are serialized by encoding some header information as elements and storing the data in lower-level structures. The fundamental structure is the *raw bitvector* that stores a binary sequence of length *n* in a vector of elements. An *integer vector* stores a sequence of *n* integers of width *w* each as a raw bitvector. Plain bitvectors store the data as a raw bitvector, followed by optional rank/select support structures. The support structures are optional and unspecified, as they are implementation-dependent and can be built quickly from the bitvector itself. Sparse bitvectors use a plain bitvector for high and an integer vector for low.

*String array* stores an ordered collection of strings as a single concatenated string, as described in Section 2.2. The serialized structure uses a sparse bitvector as an index and compresses the strings using *alphabet compaction*. For example, if the strings only contain characters in {*A, C, G, N, T*}, they are stored as an integer vector containing values in [0 … 5). In-memory structures may decompress the index as an integer vector and the strings as a byte vector for faster access.

*Dictionary* encodes a bidirectional mapping between an ordered collection of distinct strings *S*_0_, …, *S_m_*_−1_ and their identifiers [0 … *m*). In the serialized structure, the strings are stored as a string array. An integer vector stores the identifiers in lexicographic order. In-memory structures may decompress the dictionary into a more appropriate representation.

*Tags* are key-value pairs used for annotation. Keys are case-insensitive strings, and they are assumed to be distinct. Values can be arbitrary strings. Serialized tags are stored as a string array, while in-memory structures may use more appropriate representations. Key `source` identifies the library used for serializing the data. The reader may use it for determining if it can understand any optional structures that are present.

#### 2.3.3 GBWT

The serialization format for the GBWT resembles the compressed in-memory structure. It starts with a header and tags. Node records are compressed as byte sequences using the original encoding, and the sequences are concatenated in a byte vector. A sparse bitvector is used as an index over the nodes.

The GBWT uses document array samples for mapping BWT positions to the identifiers of the corresponding paths. They are serialized as an optional structure in an unspecified format. The structure is optional, because many applications do not need that functionality. The format is left unspecified, as the original structure is complex and not particularly efficient. Future versions of the file format may specify a better structure based on the r-index (Gagie *et al*., 2020).

GBWT metadata is stored in an optional structure. The structure is optional, because the GBWT may also be useful in applications outside genomics. Metadata starts with a header, followed by a vector of path names. Sample and contig names are serialized as dictionaries.

#### 2.3.4 GBWTGraph

As the GBWTGraph uses a GBWT index for graph topology, it only needs to store a header and node labels. While the in-memory data structure used in Giraffe stores the labels in both orientations for faster access, serializing the reverse orientation is clearly unnecessary. The forward labels are concatenated and stored as a string array.

While GFA graphs have segments with string names, bidirected sequence graphs have nodes with integer identifiers. And while the original graph may have segments with long labels, it often makes sense to limit the length of the labels. For example, the minimizer index used in Giraffe supports up to 1024 bp nodes. Some algorithms create temporary copies of node labels, and long labels may also be inconvenient to visualize.

For this purpose, the GBWTGraph includes a *node-to-segment translation*. The translation consists of a string array storing segment names *S*_0_, …, *S_m_*_−1_ and a sparse bitvector *B* mapping node ranges to segments. If the translation is in use, the set of nodes is assumed to be *V*′ = [1 … |*V*′|]. Segment *S_i_* is then the concatenation of nodes *v* ∈ [*B*.select(*i*, 1) … *B*.select(*i* + 1, 1)).

#### 2.3.5 GBZ

*GBZ* is a container that stores a bidirectional GBWT index and a GBWTGraph. It also includes a header and tags. The format interprets GBWT metadata in a specific way.

Paths corresponding to sample name `_gbwt_ref` are named paths. They correspond to paths in a PathHandleGraph and P-lines in GFA. The name of the path is stored as a contig name.

Other paths correspond to GFA W-lines. GBWT path names map to GFA walk metadata in the following way. Sample names and haplotype identifiers can be used directly. GBWT contig names become GFA sequence names. GBWT fragment identifier is used as the start of GFA sequence interval.

### 2.4 GFA compression

#### 2.4.1 Compression algorithm

We propose a GBZ compression algorithm that makes several passes over a memory-mapped GFA file. The algorithm works best when, for each weakly connected component in the graph and each line type, all lines from that component with that type are consecutive in the file.

In the first pass, the algorithm determines the starting position and type of each line in the file. It determines whether a node-to-segment translation is necessary. A translation is created if a segment contains a sequence longer than the user-defined threshold (1024 bp by default) or if segment names cannot be interpreted as integer identifiers. The algorithm also chooses an appropriate size for GBWT construction buffers based on the length of the longest path.

In the second pass, the algorithm processes all segments. It builds the node-to-segment translation if necessary and stores node labels in a temporary structure. The third pass processes links and creates a temporary graph. The algorithm finds the weakly connected components of the graph and sorts them by the minimum node identifier. It then determines GBWT construction jobs by using large components directly as jobs and combining consecutive small components into larger jobs.

The fourth pass builds GBWT metadata from P-lines and W-lines. If there are W-lines present, the algorithm interprets P-lines as named paths, as described in Section 2.3.5. Otherwise the algorithm breaks P-line names info fields using a user-provided regex and maps the fields to the components of GBWT path names.

GBWT construction happens in the fifth and final pass that builds a separate GBWT index for each job. The jobs are started from the largest to the smallest (by the number of nodes in the components), and multiple jobs can be run in parallel. Each job first processes P-lines and then W-lines in the corresponding components and appends the paths into a buffer. When the buffer is full, the paths are inserted into the GBWT index using the direct construction algorithm (Sirén *et al*., 2020).

After all construction jobs have finished, the algorithm merges the partial GBWT indexes into a final index. Because the jobs are based on weakly connected components, the partial indexes do not overlap, and node records from them can be used directly in the merged index (Sirén *et al*., 2020). The algorithm builds a GBWTGraph from the final GBWT index, node labels, and node-to-segment translation, and then it serializes everything in the GBZ format.

#### 2.4.2 Decompression algorithm

Decompressing GFA from a GBZ file is straightforward. We first iterate over the node-to-segment translation (or nodes if there is no translation) and output the S-lines in that order. Then we iterate over all edges in order determined by the source node, select the ones that connect segments, and output them as L-lines. Because each edge ((*v, o*), (*w, o*′)) of a bidirected sequence graph implies the existence of the reverse edge 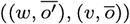, we only write the smaller of the two (in the usual tuple ordering) as an L-line. Writing L-lines is slower than writing S-lines, because it requires partial decompression of the node records. Hence it may be useful to write them using multiple threads. When multiple threads are used, we do not guarantee the exact order for performance reasons.

Then we write named paths as P-lines and finally other paths as W-lines, both according to their order in the GBWT index. We interpret GBWT metadata as described in Section 2.3.5. As extracting the paths from the GBWT index is the most expensive part of decompression, this should be done using multiple threads. Again, when multiple threads are used, we do not guarantee the exact order.

### 3 Results

#### 3.1 Experimental setup

The main GBZ library is written in C++. It builds on the original GBWT implementation. The graph supports the PathHandleGraph interface from libhandlegraph, exposing named paths through the interface. Other paths can be accessed using a custom interface. There is also a Rust library that can use existing GBWT and GBZ structures but not build new ones.

There are some differences between the implementations. The Rust implementation does not use document array samples. Because the C++ implementation uses Giraffe data structures, it decompresses and stores node labels in both orientations. The Rust implementation only stores the forward orientation. The GFA decompression algorithm in the C++ implementation uses additional memory to speed up decompression. It decompresses the node-to-segment translation and large (>1024-byte) GBWT node records for faster access. The Rust implementation uses the query interface directly without caching. Finally, the C++ implementation writes both edges and paths using multiple threads, while the Rust implementation only uses multiple threads for paths.

Both implementations use an external library for basic succinct data structures. The C++ implementation uses the vgteam fork of SDSL (Gog *et al*., 2014), which includes the sparse bitvector improvements and supports the Simple-SDS serialization format. The Rust implementation uses Simple-SDS.

As our test data, we used two 90-haplotype human graphs based on year 1 data from HPRC (Liao *et al*., 2022). Cactus was built using the Minigraph–Cactus pipeline (Hickey *et al*., 2022) with GRCh38 as the reference. PGGB was built using the Pangenome Graph Builder pipeline (Garrison *et al*., 2022). We also created a 1000GP dataset with 5008 human haplotypes by combining the full 1000 Genomes Project (The 1000 Genomes Project Consortium, 2015) indexes from the Giraffe paper (Sirén *et al*., 2021). See Table 1 for details.

**Table 1.** Datasets and their properties. We list the size of the file in uncompressed and gzip-compressed GFA format and in the GBZ format. GBWT refers to the size of the GBWT index without document array samples. We also list total sequence length (in bases) in the graph and the total length of the paths (in nodes) stored in the GBWT index, as well as the number of lines of each type in the GFA file.

| Dataset | .gfa | .gfa.gz | .gbz | GBWT | Sequence | Paths | S-lines | L-lines | P-lines | W-lines |
| --- | --- | --- | --- | --- | --- | --- | --- | --- | --- | --- |
| Cactus | 44.9 GiB | 11.1 GiB | 3.11 GiB | 1.49 GiB | 3.3 billion | 8.8 billion | 81.4 million | 113.0 million | 2580 | 24456 |
| PGGB | 88.6 GiB | 14.6 GiB | 5.73 GiB | 2.09 GiB | 8.4 billion | 16.3 billion | 110.9 million | 154.8 million | 34796 | 0 |
| 1000GP | 9534.9 GiB | 2231.3 GiB | 16.84 GiB | 7.97 GiB | 3.2 billion | 2125.1 billion | 293.2 million | 372.9 million | 0 | 115184 |

Chromosome 16 of the PGGB graph contains a highly collapsed repetitive region, where the average haplotype visits a few nodes tens of thousands or even hundreds of thousands of times. These nodes also have a large number of neighbors. This breaks the assumptions made by the GBWT, making compression much slower than expected. The C++ implementation successfully mitigates the issue during decompression by caching large node records. Hence a better in-memory representation for large records should be enough to fix this and other similar issues.

We used four systems for most experiments: Desktop (iMac 2020), Laptop (MacBook Air 2020), Intel Server (AWS i3.8xlarge), and ARM Server (AWS r6gd.8xlarge). For scalability experiments with the 1000GP dataset, we used another system with more disk space: Large (AWS i4i.16xlarge). See Table 2 for details. All data was stored on a local SSD. While we used different C++ compilers on different systems, the Rust compiler was always rustc version 1.58.1.

**Table 2.** Systems used for the experiments. Jobs indicates the number of parallel compression/decompression jobs.

| System | Processor | CPU Cores | Jobs | RAM | OS | C++ Compiler |
| --- | --- | --- | --- | --- | --- | --- |
| Desktop | Intel Core i9-10910 | 10 physical (20 logical) | 10 / 10 | 128 GiB | macOS 12.2.1 | GCC 11.2.0 |
| Laptop | Apple M1 | 4 performance + 4 efficiency | — / 4 | 16 GiB | macOS 12.2.1 | Apple Clang 13.0.0 |
| Intel Server | Intel Xeon E5-2686 v4 | 16 physical (32 logical) | 16 / 16 | 244 GiB | Ubuntu 20.04 | GCC 9.3.0 |
| ARM Server | AWS Graviton2 | 32 | 16 / 32 | 256 GiB | Ubuntu 20.04 | GCC 9.3.0 |
| Large | Intel Xeon Platinum 8375C | 32 physical (64 logical) | 32 / 32 | 512 GiB | Ubuntu 20.04 | GCC 9.3.0 |

We used the general-purpose gzip compressor to establish a baseline for performance. As gzip is often used in bioinformatics tools and pipelines, we chose it over more modern compressors such as zstd.

See Supplement 2–4 for further details.

### 3.2 Compression

We compressed the Cactus and PGGB datasets in the GBZ format using the compressor included in the C++ implementation. File sizes were 3.6 times and 2.5 times smaller, respectively, than those achieved by gzip (see Table 1). The compression ratio improves further as the number of haplotypes increases, as seen with the 1000GP dataset.

We measured the time and memory usage of GBZ compression and the time usage of gzip compression. (Memory usage of gzip compression is negligible.) The results can be seen in Table 3. Because the input GFA is memory-mapped, any parts of the input stored in disk cache are included in the memory usage of the compressor. We did not use Laptop for this benchmark due to the limited amount of memory.

**Table 3.** Wall clock time and peak memory usage for various tasks on different datasets and systems.

| Dataset | System | Compression | gzip | Loading (C++) | Loading (Rust) | Decompression (C++) | Decompression (Rust) | gunzip |
| --- | --- | --- | --- | --- | --- | --- | --- | --- |
| Cactus | Desktop | 40 min / 96.5 GiB | 25 min | 23 s / 11.8 GiB | 19 s / 5.9 GiB | 116 s / 15.5 GiB | 239 s / 7.1 GiB | 80 s |
| Cactus | Laptop | — | — | 23 s / 9.4 GiB | 16 s / 5.9 GiB | 186 s / 9.7 GiB | 304 s / 6.5 GiB | 80 s |
| Cactus | Intel Server | 19 min / 111.5 GiB | 39 min | 37 s / 11.7 GiB | 35 s / 5.9 GiB | 125 s / 14.5 GiB | 193 s / 6.5 GiB | 361 s |
| Cactus | ARM Server | 16 min / 111.0 GiB | 48 min | 33 s / 11.7 GiB | 33 s / 5.9 GiB | 86 s / 14.5 GiB | 138 s / 7.1 GiB | 350 s |
| PGGB | Desktop | 890 min / 114.7 GiB | 34 min | 43 s / 27.2 GiB | 39 s / 13.1 GiB | 309 s / 32.7 GiB | 7226 s / 14.2 GiB | 121 s |
| PGGB | Laptop | — | — | 70 s / 8.0 GiB | 40 s / 7.6 GiB | 638 s / 7.2 GiB | 18451 s / 12.5 GiB | 116 s |
| PGGB | Intel Server | 1365 min / 194.6 GiB | 54 min | 73 s / 27.1 GiB | 73 s / 13.0 GiB | 335 s / 32.2 GiB | 9194 s / 13.5 GiB | 603 s |
| PGGB | ARM Server | 1232 min / 194.3 GiB | 66 min | 65 s / 27.1 GiB | 68 s / 13.0 GiB | 187 s / 33.5 GiB | 3499 s / 14.7 GiB | 551 s |
| 1000GP | Large | 779 min / 489.2 GiB | 4341 min | 48 s / 28.1 GiB | 35 s / 12.3 GiB | 7462 s / 49.3 GiB | 7677 s / 18.8 GiB | 48618 s |

With the Cactus dataset, multi-threaded GBZ compression required similar time to single-threaded gzip compression. Intel Server and ARM Server were faster than Desktop, both with individual GBWT construction jobs and by the total running time multiplied by the number of parallel jobs. This was likely caused by the higher memory bandwidth of the server processors.

Compression speed of the PGGB dataset was dominated by the highly collapsed region of chromosome 16. Without chromosome 16, the compression would have finished in 60–90 minutes (depending on the system), which is slightly higher than gzip compression time. The GBWT construction job for chromosome 16 took an order of magnitude longer. Dealing with such collapsed regions will require changes to the in-memory data structures used in the GBWT (see Section 4). Desktop was faster than the servers due to the higher single-core speed of its processor. Its memory usage was also lower, because there was not enough memory for caching the entire input.

### 3.3 In-memory structures

The GBZ file format has been designed for fast loading into in-memory data structures. In the C++ implementation, these are the same GBWT and GBWTGraph structures that are used in the Giraffe aligner. The structures in the Rust implementation are similar. While some structures must be rebuilt when loading the file and other parts are decompressed for faster access, most of the GBWT can be copied from the file to the data structures.

We measured the time and memory usage of loading GBZ files. The results can be seen in Table 3. Loading the structures took tens of seconds. Desktop and Laptop were faster than Intel Server and ARM Server due to their higher single-core performance. The C++ implementation used more memory than the Rust implementation, because it stores node labels in both orientations.

Laptop used less memory than the other systems, especially with the PGGB dataset. This is because we defined memory usage as resident set size. When memory usage approaches memory capacity, the operating system starts swapping out inactive memory regions to compressed memory and ultimately to disk.

### 3.4 Decompression

We decompressed the Cactus and PGGB datasets from GBZ format to GFA format on all four systems using the C++ and Rust decompressors. Time and memory usage of can be seen in Table 3. We also measured the time used by gzip decompression for a comparison.

With the Cactus dataset, the multi-threaded C++ decompressor was about as fast as the single-threaded gzip decompressor on macOS. The Linux version of gzip was several times slower. The Rust decompressor was also slower, because it uses the query interface directly without caching. ARM Server was faster than Intel Server due to having more CPU cores.

Chromosome 16 in the PGGB dataset caused issues again. The C++ decompressor managed to decompress it in a reasonable time, as it caches large GBWT node records. Decompression time was reasonable even on Laptop, which only had half the memory required for the in-memory data structures. This is because the memory access patterns during decompression are mostly sequential, and swapping does not slow it down too much. The Rust implementation was more than an order of magnitude slower than the C++ implementation. Gzip decompression was again slower in Linux than in macOS.

### Scalability

We tested the scalability of the GBZ (de)compressors with the 1000GP dataset. Because the GFA file was too large for the other systems, we ran these experiments on the Large system with 13.5 TiB of local SSD space. The results can be seen in Table 3. Note that because Large is a newer system, the running times are not directly comparable with the other server instances. Loading times are comparable with Desktop, as the processors have similar single-threaded performance.

Compression took 13 hours. With the input much larger than memory capacity, roughly half of the memory usage was for in-memory data structures and half for caching parts of the input. The initial validation pass took 165 minutes, or about 1 second per gigabyte. Passes 2 to 4 (segments, links, and metadata) needed a total of 7 minutes, including 4 minutes for finding weakly connected components. The final construction pass took 605 minutes. Note that while there were enough CPU cores for 32 parallel construction jobs, there were only 23 jobs to run. Decompression times were similar with both C++ and Rust implementations, as the CPU was fast enough that disk I/O became the bottleneck.

Run-length encoded BWT is effective in compressing repeated substrings that have many copies in the file. Increasing the number of copies only increases run lengths, which take logarithmic space to encode. As haplotypes are the generally inherited in large blocks shared by many samples, the size of a GBWT index depends primarily on the number of nodes in the graph. We see this in Table 1, where the 1000GP index is only a few times larger than the Cactus and PGGB indexes despite storing over 50 times more haplotypes. We also see it in the sizes of in-memory structures, which are very similar between 1000GP and PGGB. While 1000GP needs more space for the GBWT index, the PGGB graph contains much more sequence.

See also Supplement 5 for experiments on the effect of the number of parallel jobs on compression / decompression performance.

## 4 Discussion

We have proposed the GBZ file format for pangenome graphs representing aligned genomes. The file format is based on data structures used in the Giraffe aligner, and it is the preferred graph format for the aligner. GBZ graphs are widely supported in vg (Garrison *et al*., 2018), and we also provide standalone libraries for using them in other software tools.

GBZ compresses GFA files with many similar paths well. The compression speed is competitive as long as the graph does not contain certain degenerate structures. A GBZ file can be decompressed quickly into a GFA file for tools that do not support GBZ. The graph and the paths in the file can also be loaded quickly into in-memory data structures.

There are two main data models for representing paths metadata in pangenome graphs. The W-line / GBWT model is based on structured names appropriate for assembled genomes. The P-line / libhandlegraph model uses string names, which are harder to interpret but suitable for more diverse applications. GBZ supports both models. Some applications of pangenome graphs require specifying a set of paths that acts as a linear reference sequence. Neither model has a way of specifying whether the graph contains a preferred reference sequence or if any sample is an equally valid choice. If such a convention emerges, GBZ can support it using tags.

The GBWT assumes that the nodes of the graph do not have too many neighbors and the paths do not visit the same nodes too many times. Pangenome graph construction pipelines sometimes produce graphs that violate these assumptions. Because the GBWT has to decompress the node record every time it computes LF-mapping from that node, this can cause significant performance loss. However, as we saw in the decompression benchmarks (Section 3.4), caching large records in a more suitable format helps to avoid the issue. While the format we used was only appropriate for extracting paths from the GBWT, a proper solution such as a run-length encoded wavelet tree should work in most situations. The ones from the DYNAMIC library (Prezza, 2017) look promising. However, using them in the GBWT may require significant engineering effort and changes to the query interface.

## Acknowledgements

We would like to thank members of the Human Pangenome Reference Consortium, in particular Glenn Hickey, Heng Li and Erik Garrison for providing pangenome graphs to experiment with. We also thank Adam M. Novak for the idea of a combined GBWT + GBWTGraph file format and for contributions to the C++ implementation. Finally, we would like to thank everybody in the VG Team.

## Funding

This work was supported by the National Human Genome Research Institute of the National Institutes of Health [U01HG010961, R01HG010485, U41HG010972, U24HG011853, OT2OD026682]. The content is solely the responsibility of the authors and does not necessarily represent the official views of the National Institutes of Health.

## Footnotes

1 The software versions used in this paper output GFA 1.0, as the status of W-lines in the specification was still unclear when we started the (de)compression benchmarks.

## Notes

https://github.com/jltsiren/gbwtgraph

https://github.com/jltsiren/gbwt-rs

### Competing Interest Statement

The authors have declared no competing interest.

### Summary of Updates

Compression / decompression results with the 1000GP graph. Text clarifications. A figure giving and overview of the file format. Scaling with the number of compression / decompression jobs (in the supplement).

