## Supplementary Material for "GBZ File Format for Pangenome Graphs"

September 6, 2022

#### 1 File formats

##### 1.1 Simple-SDS

GBZ file format is based on Simple-SDS serialization, which is defined at `https://github.com/jltsiren/simple-sds/blob/main/SERIALIZATION.md`. We used version 0.2 of the serialization format. The document describes the basic principles for serialization. It also includes serialization formats for vectors, strings, bit-packed integer vectors, and plain and sparse bitvectors.

##### 1.2 GBWT

The Simple-SDS serialization format for GBWT is defined at `https://github.com/jltsiren/gbwt/blob/master/SERIALIZATION.md`. We used GBWT file format version 5 and metadata file format version 2. The document also defines serialization for string arrays, dictionaries, and tags.

##### 1.3 GBZ

The GBZ file format itself is defined at `https://github.com/jltsiren/gbwtgraph/blob/master/SERIALIZATION.md`. We used version 1 of the GBZ file format and version 3 of the GBWTGraph file format.

#### 2 Software

##### 2.1 C++ implementation

The C++ implementation is available at `https://github.com/jltsiren/gbwtgraph`. We used version 0.8.1. Its dependencies include:

- GBWT (`https://github.com/jltsiren/gbwt`) version 1.3.1;
- vgteam fork of SDSL (`https://github.com/vgteam/sdsl-lite`) version 2.3.1; and

- libhandlegraph (`https://github.com/vgteam/libhandlegraph`), git commit `21556f1e23720d7404d2478bf43fce772fe1151f` (2022-01-03).

We compiled it using the included makefile.

##### 2.2 Rust implementation

The Rust implementation can be found at `https://github.com/jltsiren/gbwt-rs`. We used versions 0.2.1 and 0.2.2. The only difference between the versions is that 0.2.2 allows using more decompression threads. The implementation depends on:

- Simple-SDS (`https://github.com/jltsiren/simple-sds`) version 0.3.1;
- getopts (`https://crates.io/crates/getopts`) version 0.2;
- libc (`https://crates.io/crates/libc`) version 0.2;
- Rand (`https://crates.io/crates/rand`) version 0.8; and
- Rayon (`https://crates.io/crates/rayon`) version 1.5.

We compiled it in release mode with `cargo build --release --features=binaries`.

#### 3 Datasets

##### 3.1 HPRC graphs

We used graphs based on year 1 data from the Human Pangenome Reference Consortium for compression and decompression benchmarks. The graphs and some information about them can be found at `https://github.com/human-pangenomics/hpp_pangenome_resources/`.

Cactus graph was built using the Minigraph–Cactus pipeline with GRCh38 as the reference. It can be downloaded from
`https://s3-us-west-2.amazonaws.com/human-pangenomics/pangenomes/freeze/freeze1/minigraph-cactus/hprc-v1.0-mc-grch38.gfa.gz`.

PGGB graph was built using the Pangenome Graph Builder pipeline. It can be downloaded from `https://s3-us-west-2.amazonaws.com/human-pangenomics/pangenomes/freeze/freeze1/pggb/hprc-v1.0-pggb.gfa.gz`.

##### 3.2 1000 Genomes Project graph

The 1000GP graph is the full unfiltered 1000 Genomes Project graph used in the Giraffe paper. We downloaded the existing GBWT index from
`https://cgl.gi.ucsc.edu/data/giraffe/mapping/graphs/for-NA19239/1000gplons/hs38d1/1000GPlons_hs38d1.gbwt`
and the XG graph from

https://cgl.gi.ucsc.edu/data/giraffe/mapping/graphs/for-NA19239/1000gplons/hs38d1/1000GPlons_hs38d1.xg

We converted the files into a GBZ graph using VG version 1.38.0 and the following command:

```
vg gbwt -x $GRAPH.xg -g $GRAPH.gbz --gbz-format $GRAPH.gbwt
```

This reads an XG graph `$GRAPH.xg` and a GBWT index `$GRAPH.gbwt` and outputs a GBZ graph `$GRAPH.gbz`. The options are:

- `-x <FILE>`: Load a graph from `<FILE>` in any supported format.
- `-g <FILE>`: Build a GBWTGraph and write it to `<FILE>`.
- `--gbz-format`: Serialize the GBWTGraph in the GBZ format instead of using separate files for the graph and the GBWT index.

### 4 Benchmarks

Before running each command, we flushed disk caches. This was done using `sync && sudo purge` in macOS and `sync && echo 3 | sudo tee /proc/sys/vm/drop_caches` in Linux.

We measured running time (wall clock time) and memory usage (maximum resident set size). Our tools measure them internally. With other tools, we used `/usr/bin/time -l` in macOS and `/usr/bin/time -v` in Linux.

#### 4.1 GBZ compression

We used the `gfa2gbwt` binary included in the C++ implementation with the following arguments:

```
gfa2gbwt -p -P $JOBS $GRAPH
```

This reads a GFA file `$GRAPH.gfa` and writes a GBZ file `$GRAPH.gbz`. The options are:

- `-p`: Show progress information, including running time and memory usage.
- `-P <INT>`: Run `<INT>` GBWT construction jobs in parallel.

The PGGB graph stores all paths as GFA P-lines. In order to parse GBWT metadata from GFA path names, we used the following options:

- `-r "(.*)#(.*)"`: Separate the path name into two fields at the first occurrence of `#`.
- `-f XSC`: Ignore the full path name (`X`), use the first field as a sample name (`S`), and use the second field as a contig name (`C`).

##### 4.2 GBZ loading, C++

We used the gfa2gbwt binary included in the C++ implementation with the following arguments:

```
gfa2gbwt -l -p $GRAPH
```

This reads a GBZ file $GRAPH.gbz. The options are:

- -l: Load GBZ, no output (hidden option).
- -p: Show progress information, including running time and memory usage.

##### 4.3 GBZ loading, Rust

We used the gbunzip binary included in the Rust implementation with the following arguments:

```
gbunzip -l -v $GRAPH.gbz
```

This reads a GBZ file $GRAPH.gbz. The options are:

- -l: Load GBZ, no output.
- -v: Show progress information, including running time and memory usage.

##### 4.4 GBZ decompression, C++

We used the gfa2gbwt binary included in the C++ implementation with the following arguments:

```
gfa2gbwt -d -p -P $JOBS $GRAPH
```

This reads a GBZ file $GRAPH.gbz and writes a GFA file $GRAPH.gfa. The options are:

- -d: Decompress GFA from GBZ.
- -p: Show progress information, including running time and memory usage.
- -P <INT>: Run <INT> decompression threads in parallel.

##### 4.5 GBZ decompression, Rust

We used the gbunzip binary included in the Rust implementation with the following arguments:

```
gbunzip -v -t $JOBS $GRAPH.gbz > $GRAPH.gfa
```

This reads a GBZ file $GRAPH.gbz and writes a GFA file $GRAPH.gfa. The options are:

- -v: Show progress information, including running time and memory usage.
- -t <INT>: Run <INT> decompression threads in parallel.

##### 4.6 Comparison with gzip

We compared the compression and decompression speed of our tools to gzip. For compression, we used the following command:

```
gzip -c $GRAPH.gfa > $GRAPH.gfa.gz
```

This reads a GFA file `$GRAPH.gfa` and compresses it to `$GRAPH.gfa.gz` with the default compression level.

For decompression, we used:

```
gunzip -c $GRAPH.gfa.gz > $GRAPH.gfa
```

This reads a gzip file `$GRAPH.gfa.gz` and decompresses it to `$GRAPH.gfa`.

#### 5 Number of (de)compression jobs

We measured the scaling of GBZ compression and decompression by varying the number of parallel jobs. We restricted our attention to the C++ implementation, the well-behaving Cactus dataset, and the ARM Server and Intel Server systems. The results can be seen in Figure 1.

In compression, only the final construction pass is parallelized. With 16 parallel jobs, the sequential parts took 43 % and 44 % of total running time on ARM Server and Intel Server, respectively. (For a comparison, the sequential parts took only 22 % of the total with the 1000GP dataset, where the total length of the paths is much larger.) We ultimately get 6x and 5x scaling with 14 parallel jobs on ARM Server and Intel Server, respectively. Increasing the number of parallel jobs beyond 14 does not help anymore, as the largest individual jobs (which generally correspond to chromosomes) become the bottleneck.

There are two bottlenecks in decompression: disk bandwidth and loading the GBZ file into in-memory data structures. On ARM Server, which has only a single SSD, we start seeing diminishing returns after 12 parallel jobs, as we have saturated disk bandwidth. Intel Server, which has four SSDs, continues to scale until we run out of physical CPU cores. We achieved 7x scaling on both systems. While we did not measure the scaling of the Rust decompressor, we expect that it will need more parallel jobs to saturate disk bandwidth. As we saw with the 1000GP dataset in Section 3.5, if there are enough CPU cores, both decompressors will ultimately achieve similar speed.

We also measured the time/space trade-offs achieved by varying the number of parallel construction jobs. (In decompression, the number of parallel jobs has a negligible impact on memory usage.) The results can be seen in Figure 2. As the number of parallel jobs increases, memory usage increases from 78 GiB to 111 GiB. This becomes 33 GiB to 66 GiB if we ignore the cached input. Which, as we saw in Section 3.5, does not have a significant impact on compression speed. Hence the trade-offs achieved with 4 to 8 parallel jobs are attractive on memory-constrained systems.

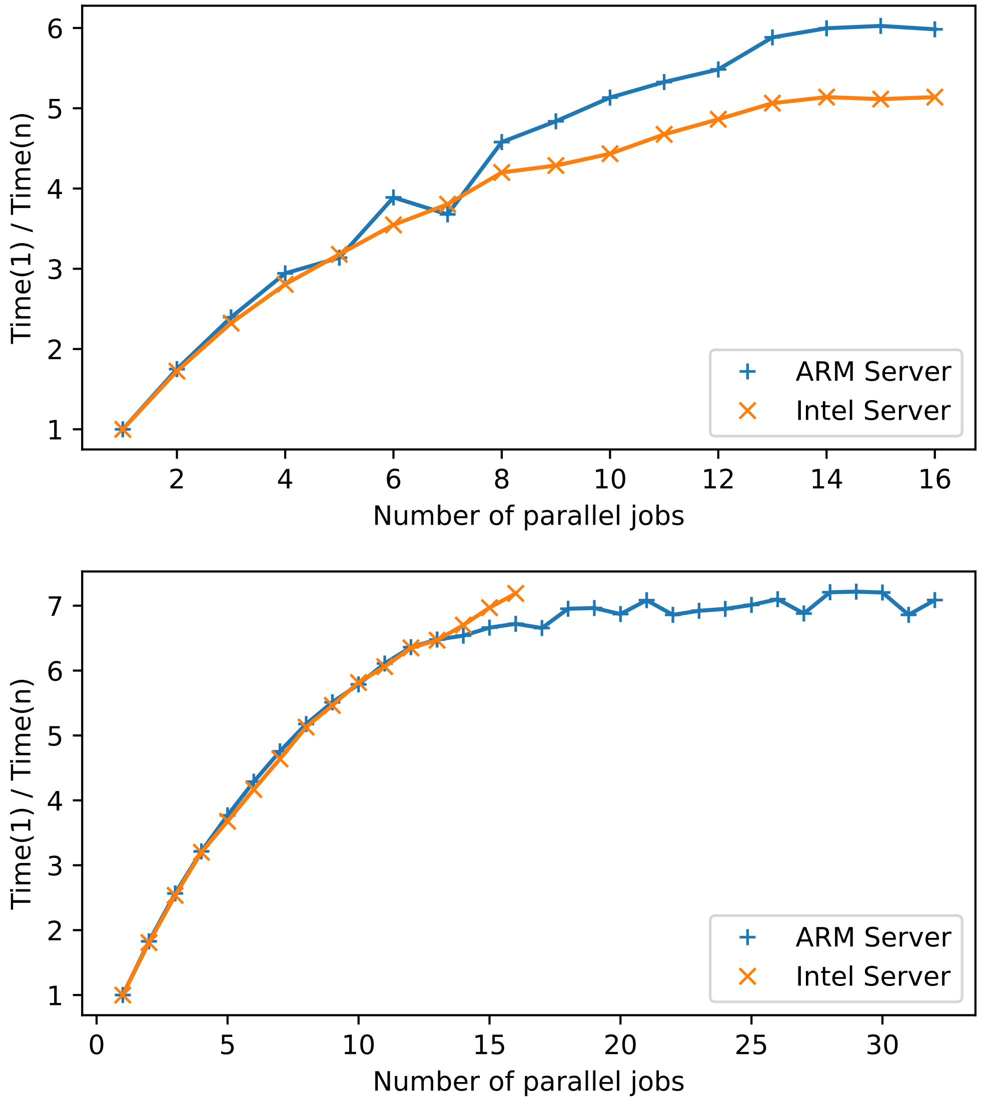


Figure 1: Scaling of GBZ compression (top) and decompression (bottom) with the number of parallel jobs. The vertical axis is the running time with a single job divided by the running time with $n$ parallel jobs.

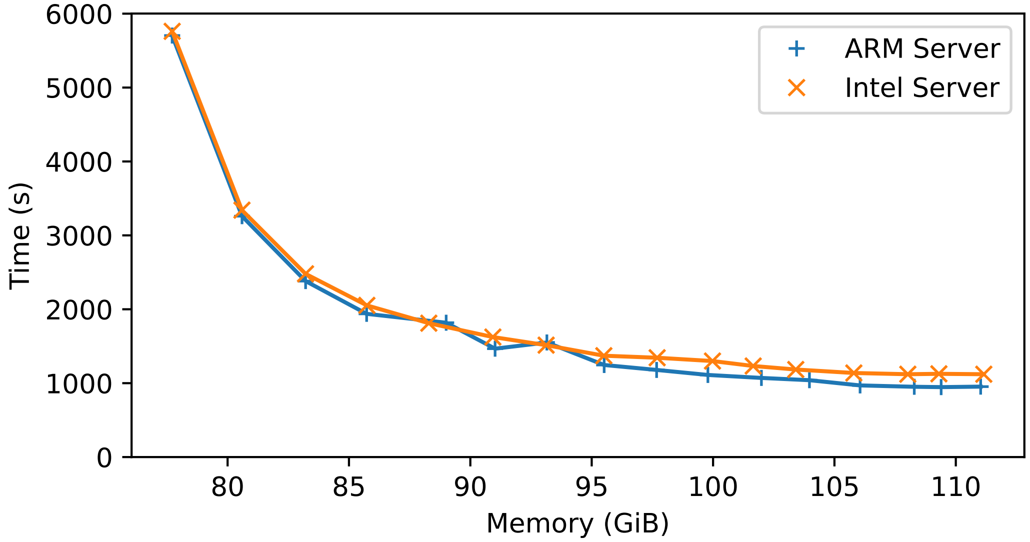


Figure 2: Time/space trade-offs for GBZ compression by varying the number of parallel construction jobs.
